## Supplementary material for "Archaic introgression contributed to shape the adaptive modulation of angiogenesis and cardiovascular traits in human high-altitude populations from the Himalayas": Figure 1-4 - figure supplement

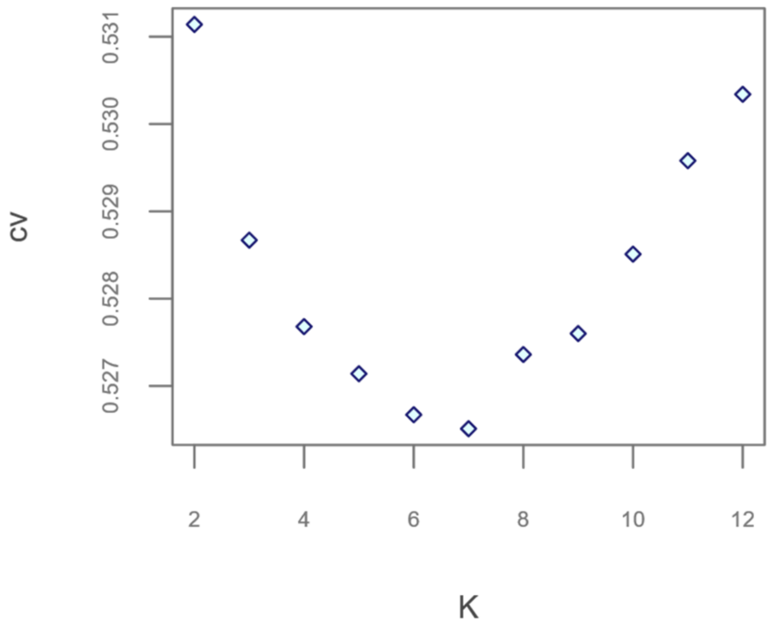


**Figure 1 - figure supplement 1.** Scatterplot showing the number of possible population clusters (K) tested by the different ADMIXTURE runs performed and the Cross Validation (CV) errors associated to them.

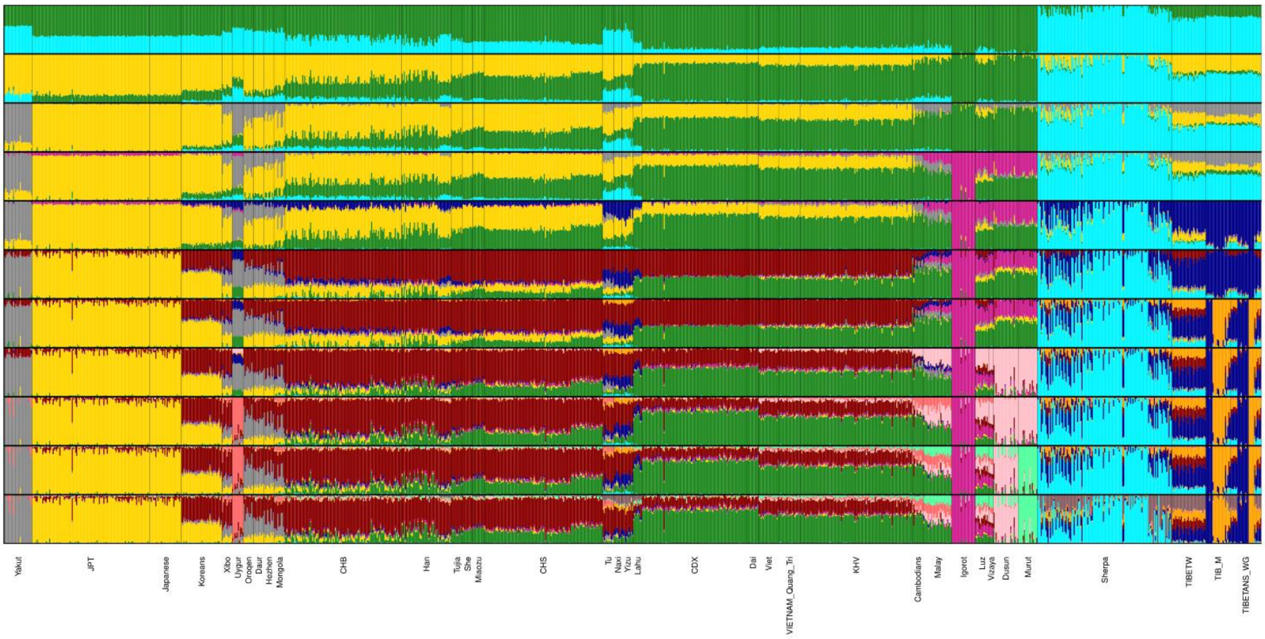


**Figure 1 – figure supplement 2.** Admixture analyses performed on the extended dataset for K = 2 to K = 12. Numbers of K increase from the top to the bottom of the plot and the considered populations are named according to the labels reported in Supplement table 1.

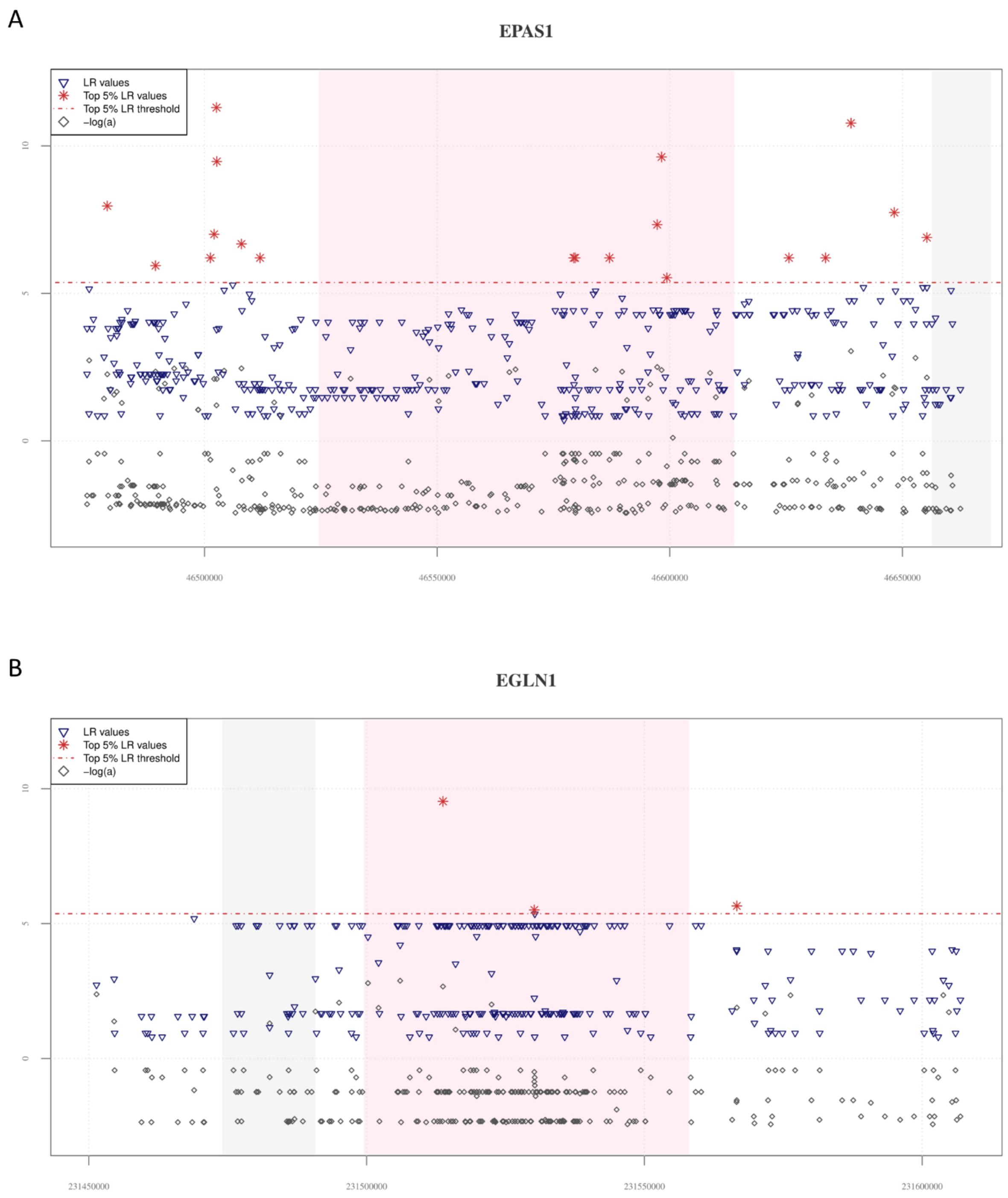


**Figure 2 – figure supplement 1.** Distribution of *VolcanoFinder* statistics across the *EPAS1* and *EGLN1* positive and negative controls for adaptive introgression. In the plots, the chart base reports genomic positions of variants, pink background indicates the starting and the ending positions of the genes, while the grey background identifies those genes (i.e., the *LINC02583* gene in the *EPAS1* downstream genomic region, the *SPRTN* gene in the upstream *EGLN1* region) possibly involved in regulatory transcription mechanisms. The dashed red line identifies the significant threshold set to filter likelihood ratio values (i.e., top 5% LR values). **(A)** The 19 SNVs showing significant likelihood ratio values (i.e., red stars) resulted closely distributed in the ending portion of the *EPAS1* gene and in both up and downstream genomic regions flanking such locus. The -logα values (i.e., grey diamonds) appeared consistently distributed in the entire *EPAS1* region. A similar a pattern is observed also for the *TBC1D1*, *PRKAG2* and *RASGRF2* new candidate AI genes, as reported in Figure 2A and 2B and in Figure 2 – figure supplement 3. **(B)** The *EGLN1* genomic region is characterized by only three LR significant values among which only one strongly deviates from the significant LR threshold. Several SNVs distributed in the *EGLN1* starting portion, as well as in its flaking regions, showed elevated -logα values supporting the action of natural selection on them in the considered Tibetan population.

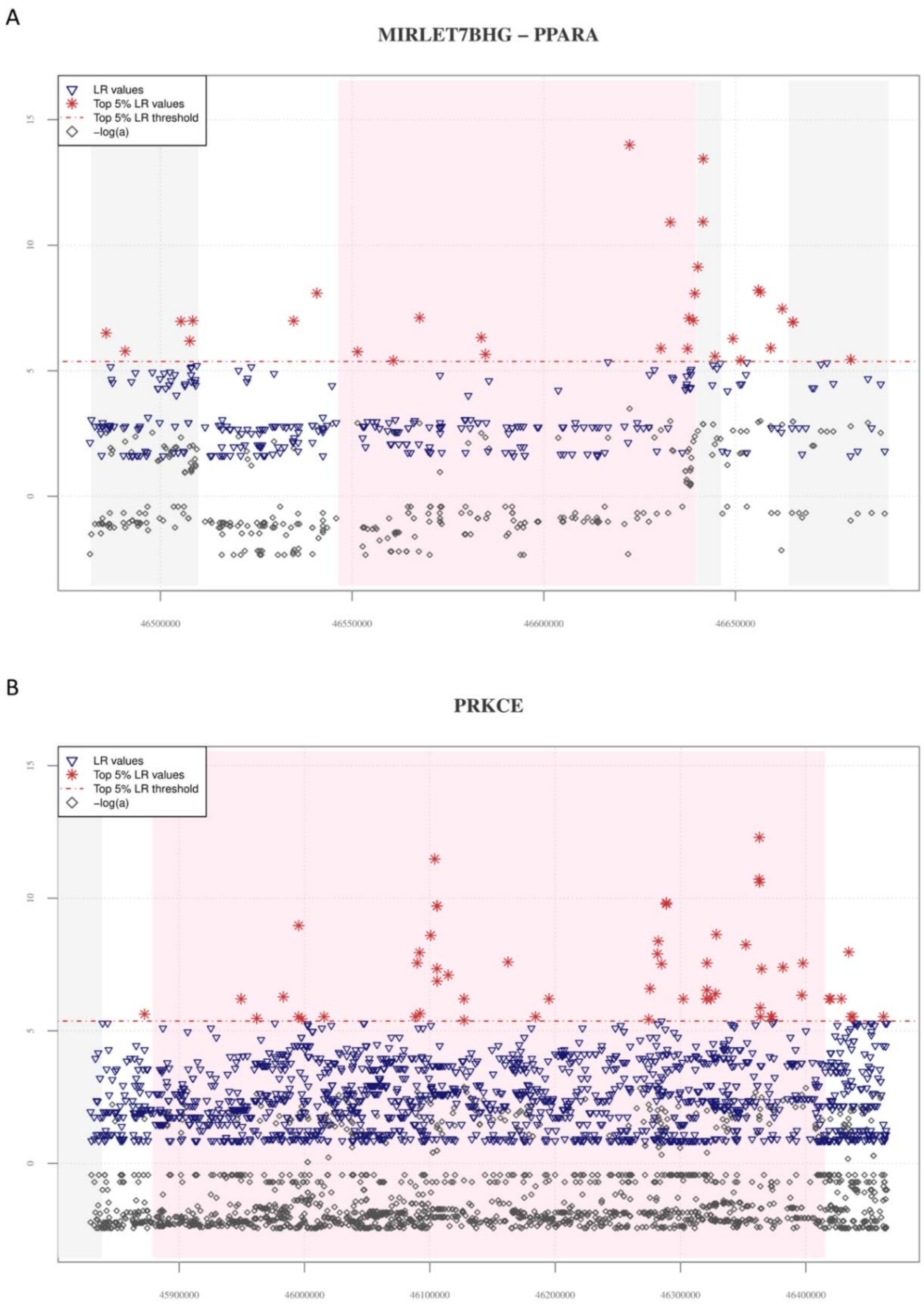


**Figure 2 – figure supplement 2.** Distribution of *VolcanoFinder* statistics across *MIRLET7BHG*, *PPARA* and *PRKCE* genes. In the plots, the chart base reports genomic positions of variants, pink background indicates the starting and the ending positions of the *PPARA* and *PRKCE* genes, while the grey background identifies those loci (i.e., *MIRLET7BHG* long non-coding, *CDPF1* and *TTC38* genes located in *PPARA* upstream and downstream regions and the *SRBD1* gene located in the *PRKCE* upstream genomic region, respectively) possibly involved in regulatory transcription mechanisms. The red horizontal dashed line displayed the significant threshold set for filtering LR significant values. **(A)** A total of 32 SNVs showed significant LR values (red stars) covering all the genomic region considered. LR greatest peaks are observed in the ending portion of the *PPARA* gene and in the *CDPF1* gene. Collectively, the genomic regions comprising significant LR scores also showed elevated -logα values (grey diamonds). **(B)** The 55 significant LR scores and the elevated -logα values cover the entire region of the *PRKCE* gene (i.e., a gene located in a genomic region nearby to *EPAS1*).

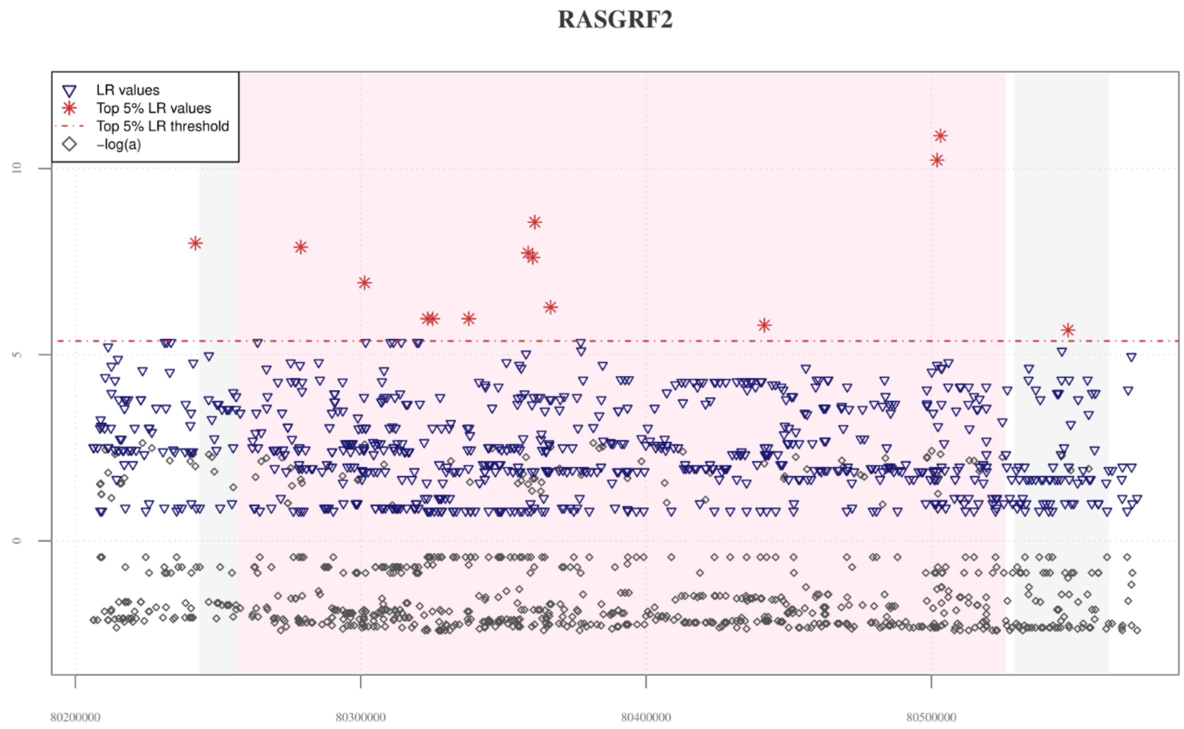


**Figure 2 – figure supplement 3.** Distribution of *VolcanoFinder* statistics across the *RASGRF2* candidate AI gene. The chart base reports genomic positions of variants, pink background indicates the starting and the ending positions of the *RASGRF2* gene, while the grey background identifies those genes (i.e., *RASGRF2-AS1* antisense RNA gene and *CKMT2* located in the *RASGRF2* upstream and downstream regions, respectively) possibly involved in regulatory transcription mechanisms. The red horizontal dashed line displayed the significant threshold set for filtering LR significant values. A total of 14 significant LR values (red stars) were equally distributed across the entire genomic region considered, with the greatest peak recovered in the ending portion of the *RASGRF2* gene. Elevated peaks of -logα are observable in both *RASGRF2* gene and its flanking genomic regions. Such a pattern resulted in line with that observed for the *EPAS1* positive control for AI, as reported in Figure 2 - figure supplement 1, and for both *TBC1D1* and *PRKAG2* genomic regions, as reported in Figure 2A and 2B.

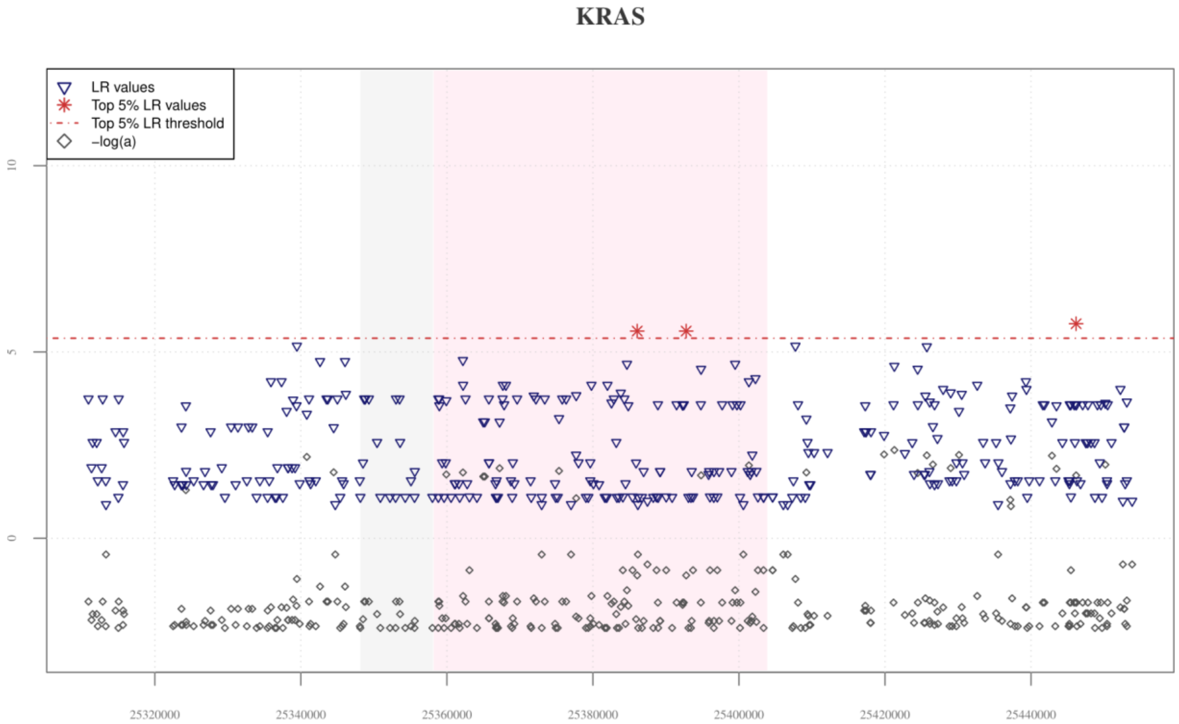


**Figure 2 – figure supplement 4.** Distribution of *VolcanoFinder* statistics across the *KRAS* candidate AI gene. The pink and the grey rectangulars represent the portion of the genome covered by the *KRAS* and *ETFRF1* genes, respectively. Although only three significant LR values can be observed for such a genomic region, the *KRAS* gene was included in our set of new candidate AI genes because it was confirmed by all the subsequent validation analyses performed and according with previous evidenced advanced by Hu et al. (2017) and Browning et al. (2018), which suggest that a portion of variants included in the *KRAS* gene, as well as in its downstream region, shows signatures of archaic Denisovan introgression in both Tibetan and CHB populations. Elevated -logα values are instead consistently distributed across the *KRAS* gene and its surrounding genomic regions.

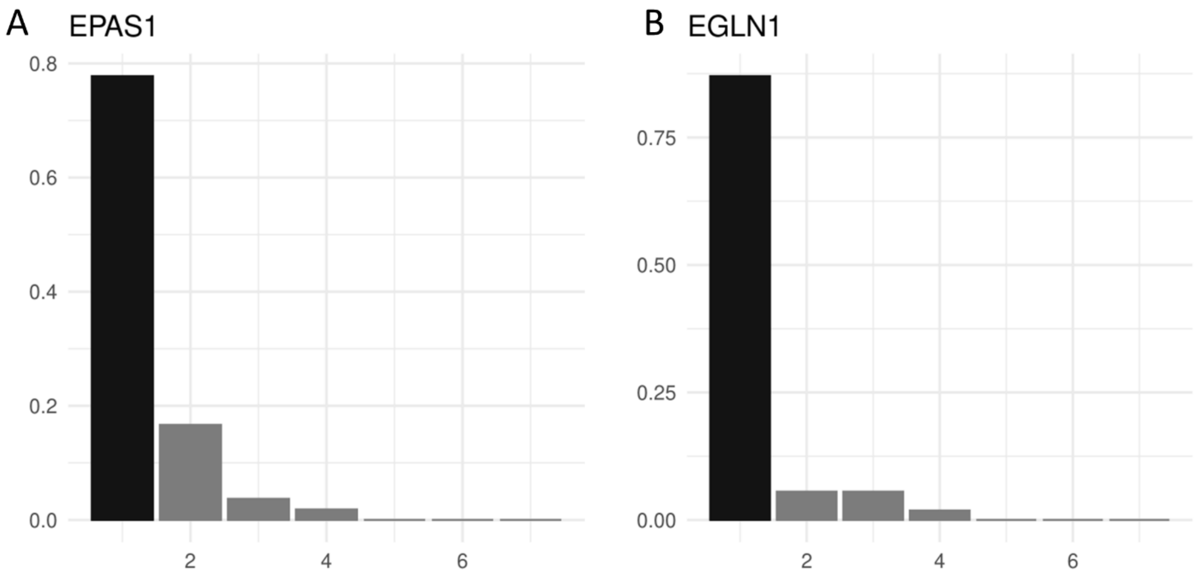


**Figure 4 – figure supplement 1.** Haplotype frequency spectra of the top windows detected as adaptively evolved by *LASSI* in the *EPAS1* and *EGLN1* genomic regions. Barplots showing haplotype frequency spectra in the genomic windows associated with the highest *T* value and linked to **(A)** *EPAS1* and **(B)** *EGLN1* genes. The x axis reports the haplotypes detected in the windows, while on y axes are indicated frequencies of each haplotype. For these windows the haplotype frequency spectra clearly reflect the pattern of diversity expected under the hard selective sweeps model in which a single predominant haplotype carrying adaptive variants (i.e., sweeping haplotypes represented with the black bars) reaches elevated frequencies in the population.

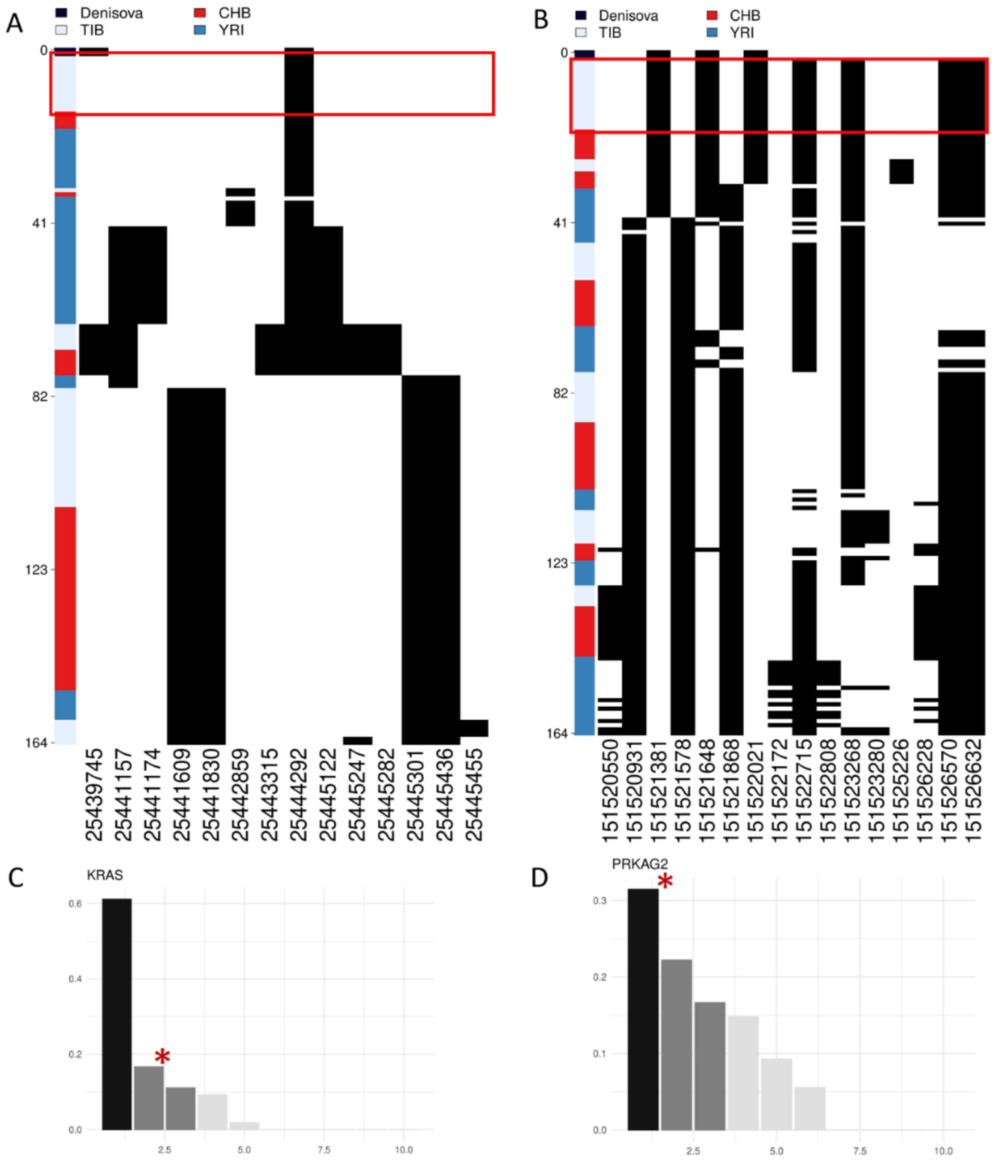


**Figure 4 - figure supplement 2.** Representation of genetic distances between modern and archaic haplotypes. Heatmap displaying the divergence between Tibetan, CHB and YRI *KRAS* and PRKAG2 haplotypes with respect to the Denisovan sequence. Haplotypes are reported in rows, while derived (i.e., black square) and ancestral (i.e., white square) alleles are displayed in columns. Haplotypes are ranked from top to bottom according to their number of pairwise differences with respect to the Denisovan sequence. The red square identifies the cluster of Tibetan haplotypes classiefed by the *LASSI* method as sweeping haplotypes (i.e., haplotypes with elevated or moderate frequencies and which carry putative adaptive variants). **(A)** 16% of Tibetan haplotypes inferred for *KRAS* conformed with a non-neutral evolutionary scenario according to *LASSI* results and presented the smallest number of pairwise difference with respect to the Denisovan genome, being plotted in the upper part of the heatmap. **(B)** 33% of Tibetan *PRKAG2* haplotypes cluster in the upper part of the heatmap being among the most close haplotypes with respect to the Denisovan sequence and presenting only four pairwise differences with it. Barplots showing haplotype frequency spectrum of *KRAS* and *PRKAG2* windows suggestive of adaptations mediated by soft selective sweeps in Tibetans. In both the plots are reported on the x axis the haplotypes detected in the considered windows, while on y axes are indicated the frequencies of each haplotype. The black and dark-grey bars indicate the more frequent haplotypes (i.e., the sweeping haplotypes inferred by *LASSI*), while the red star marks those haplotypes carrying Denisovan-like derived variants. **(C)** *KRAS* presents a pattern qualitatively comparable to that expected for a non-neutral evolution (i.e, positive likelihood T values), with two main haplotypes carrying putative adaptive variants and reaching elevated frequencies in Tibetans. The second sweeping haplotype carries the Denisovan-like derived variant and reaches 16% of frequency. **(D)** The most frequent sweeping haplotype detected in this *PRKAG2* window reches 33% of frequency in Tibetans and carries the Denisovan-like derived variant.

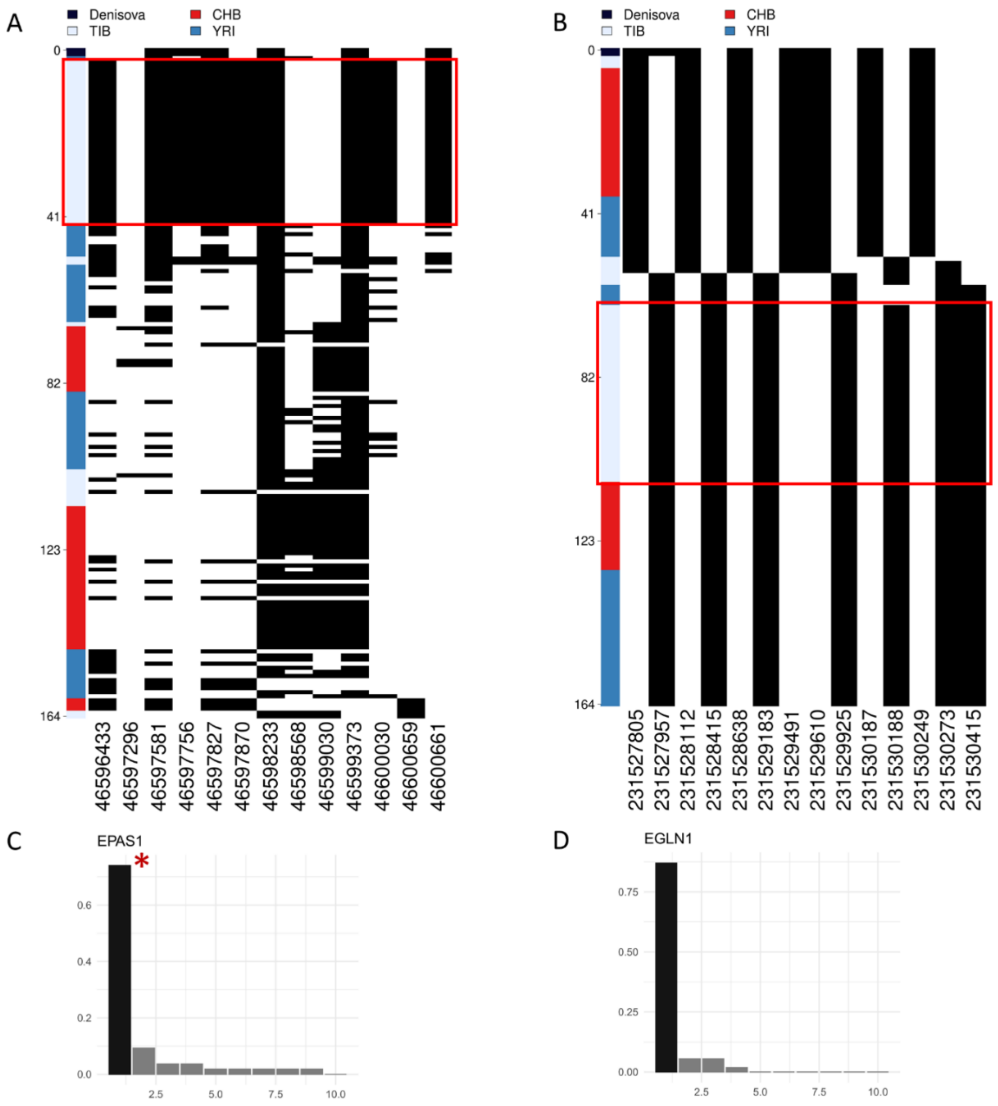


**Figure 4 - figure supplement 3.** Representation of genetic distances between modern and archaic haplotypes. Heatmap displaying the divergence between Tibetan, CHB and YRI *EPAS1* and *EGLN1* haplotypes with respect to the Denisovan sequence. Haplotypes are reported in rows, while derived (i.e., black square) and ancestral (i.e., white square) alleles are displayed in columns. Haplotypes are ranked from top to bottom according to their number of pairwise differences with respect to the Denisovan sequence. The red square identifies the cluster of Tibetan haplotypes classified by *LASSI* as sweeping haplotypes (i.e., haplotypes with elevated or moderate frequencues which carry putative adaptive variants). **(A)** The first homogeneus cluster of haplotypes visible in upper part of the heatmap belongs to Tibetan individuals (i.e., light-blue cluster). These haplotypes are among the closest ones to the archaic Denisovan sequence indicated in black, thus confirming archaic introgression at *EPAS1*. As concerning the haplotypes inferred for the other population in the plot, only one YRI haplotype presents one pairwise difference less than the haplotypes in the first Tibetan cluster. **(B)** Except for two Tibetan haplotypes, which appeared the closest ones to the archaic reference, the cluster of haplotypes presenting the lowest number of differences with respect to the Denisovan sequence belongs to the Han Chinese population. The most frequent Tibetan haplotype did not present any variant shared with the archaic reference, counting 14 pairwise differences with respect to Denisovan genome and thus not supporting the archaic origin of these *EGLN1* variants. Barplots showing the haplotype frequencies spectrum of the *EPAS1* and *EGLN1* windows suggestive of adaptation mediated by hard selective sweeps in Tibetans. In both the plots are reported on the x axis the haplotypes detected in the considered windows, while on y axes are indicated the frequencies of each haplotype. The black and dark-grey bars indicate the more frequent haplotypes (i.e., sweeping haplotypes inferred by *LASSI*), while the red star marks those sweeping haplotypes carrying Denisovan-like derived variants. For these windows, haplotype frequency spectra clearly reflect patterns expected under the hard selective sweep model in which a single haplotype carrying adaptive variants (i.e., sweeping haplotypes represented with the black bars) reaches elevated frequencies in the population. However, only for *EPAS1* such haplotype effetively carries the Denisovan-like derived variant.

### References

Browning, S. R., Browning, B. L., Zhou, Y., Tucci, S., & Akey, J. M. (2018). Analysis of Human Sequence Data Reveals Two Pulses of Archaic Denisovan Admixture. *Cell*, *173*(1), 53-61.e9. https://doi.org/10.1016/j.cell.2018.02.031

Harris, A. M., & DeGiorgio, M. (2020). A likelihood approach for uncovering selective sweep signatures from haplotype data. *Molecular Biology and Evolution*, *37*(10), 3023–3046. https://doi.org/10.1093/molbev/msaa115

Hu, H., Petousi, N., Glusman, G., Yu, Y., Bohlender, R., Tashi, T., Downie, J. M., Roach, J. C., Cole, A. M., Lorenzo, F. R., Rogers, A. R., Brunkow, M. E., Cavalleri, G., Hood, L., Alpatty, S. M., Prchal, J. T., Jorde, L. B., Robbins, P. A., Simonson, T. S., & Huff, C. D. (2017). Evolutionary history of Tibetans inferred from whole-genome sequencing. *PLoS Genetics*, *13*(4). https://doi.org/10.1371/journal.pgen.1006675
