## Supplement information for "Archaic introgression contributed to shape the adaptive modulation of angiogenesis and cardiovascular traits in human high-altitude populations from the Himalayas"

### Materials and methods

#### *VolcanoFinder* algorithm

The *VolcanoFinder* algorithm (Setter et al., 2020) was used to carry out a scan of the modern genomes, by calculating two values: α (subsequently converted into -logα) and the composite Likelihood Ratio (LR) statistic, which are respectively informative of the selection strength and of the conformity to the evolutionary model of adaptive introgression predicted for each polymorphic site. Such model considers three populations: recipient (i.e. the modern population), donor (i.e. the archaic population), and outgroup and assumes the occurrence of the introgression event after which a beneficial haplotype is introduced in the modern gene pool and starts to rise in frequency because of demographic random processes and because of the action of natural selection on it. The influence of these evolutionary forces comports the existence of multiple haplotypes that carry the beneficial archaic allele, thus comporting elevated heterozygosity level tested by the LR statistic. Therefore, the pattern of variability tested by such a statistic deeply differs from that attributable to classical selective sweeps (especially from that associated to a hard selective sweep), allowing to identify weak signatures of adaptation that resulted from both the introgression and the action of natural selection on beneficial standing genetic variation.

#### *Signet* algorithm

We first assessed the archaic state of all variants included in our dataset by comparing Tibetan genomes with the archaic Denisovan sequence, by keeping only those variants in common between them. After that, we decided to compare the set of the resulted archaic alleles with the ancestral reconstructed refence human genome sequence in order to infer the ancestral or derived state of each allele. According with this procedure, we were able to keep only those alleles that were characterized by both an archaic and a derived state thus considering simultaneously those mutations in common between Tibetans and Denisovans. Moreover, we filtered these common alleles also in function of an outgroup population (i.e., Yorubas from the 1000 Genomes Project panel of African populations), by keeping only those Denisovan-like derived alleles that have frequency equal to zero in the Yoruba population. We then associated to each gene the number and the average frequency of archaic/derived alleles, building the input file necessary for *Signet*. The *Signet* approach consists in crosschecking the information contained in the input dataset whit that collected in reference databases of functional pathways, such as the Kyoto Encyclopaedia of Genes and Genomes (KEGG) (available at: https://www.kegg.jp/), using a simulated annealing algorithm approach to define the High Scoring Subnetworks (HSS) within each biological pathway (Gouy & Excoffier, 2020). Using this approach allowed us to indagate the impact of Denisovan archaic introgression on specific biological functions in order to validate results obtained with the *VolcanoFinder* method.

#### *LASSI* algorithm

The rationale behind the *LASSI* approach (Harris & DeGiorgio, 2020) is based on the recognition of the modification resulted in the haplotype frequency spectrum of a given genomic region after the action of natural selection on it. According to the hard sweep model, when in the population arise a beneficial mutation with a very strong impact on a given phenotypic trait, the haplotype frequency spectrum of such a genomic region will be characterized by a single haplotype with an extremely elevated frequency in the population (i.e., the sweeping haplotype), while the other haplotypes whether they exist, are found at very low frequencies. Consequentially, when the selection acts on few new variants with a lower impact on a trait or on standing genetic variation, the resulted haplotype spectrum will be characterized by the existence of two or few more haplotypes that reach moderate frequencies in the population. Conversely, the variability pattern associated to the haplotype frequency spectrum expected under neutrality will be characterized by a series of different haplotypes at low frequencies in the population.

### References

Gouy, A., & Excoffier, L. (2020). Polygenic Patterns of Adaptive Introgression in Modern Humans Are Mainly Shaped by Response to Pathogens. *Molecular Biology and Evolution*, *37*(5), 1420–1433. https://doi.org/10.1093/molbev/msz306

Harris, A. M., & DeGiorgio, M. (2020). A likelihood approach for uncovering selective sweep signatures from haplotype data. *Molecular Biology and Evolution*, *37*(10), 3023–3046. https://doi.org/10.1093/molbev/msaa115

Setter, D., Mousset, S., Cheng, X., Nielsen, R., DeGiorgio, M., & Hermisson, J. (2020). VolcanoFinder: Genomic scans for adaptive introgression. In *PLoS Genetics* (Vol. 16, Issue 6). https://doi.org/10.1371/journal.pgen.1008867
